# Neural mechanism of acute stress management by trace aminergic signalling in the lateral habenula

**DOI:** 10.1101/2022.02.02.478114

**Authors:** Soo Hyun Yang, Esther Yang, Jaekwang Lee, Jin Yong Kim, Hyeijung Yoo, Hyung Sun Park, Jin Taek Jung, Dongmin Lee, Sungkun Chun, Hyun Woo Lee, Hyun Kim

**Affiliations:** Department of Anatomy, College of Medicine, Korea University, Seoul 02841, Korea; Division of Functional Food Research, Korea Food Research Institute, Wanju, 55365, Korea; Department of Physiology, Jeonbuk National University Medical School, Jeonju 54907, Korea

## Abstract

Stress management is a prerequisite for the survival of vertebrates because chronic stress may cause depression and is known to negatively modulate the dopaminergic reward system^1^. Enhanced excitability of neurons in the lateral habenula (LHb) induced by chronic stress is essential for silencing dopaminergic neurons in the ventral tegmental area (VTA) via GABAergic neurons in the rostromedial tegmental nucleus (RMTg)^2–10^. However, the effect of acute stress on the LHb-RMTg-VTA pathway is unknown^11–14^. Here, we show that both aromatic L-amino acid decarboxylase (AADC)-expressing neurons (D-neurons)^15^ in the LHb and dopaminergic neurons in the VTA are activated by acute stress, whereas GABAergic neurons in the RMTg are not. Selective stimulation of LHb D-neurons and AADC knockdown of these neurons reverse-regulate the RMTg-VTA pathway. Circuit tracing and electrophysiology data demonstrate that trace aminergic signalling by LHb D-neurons directly suppresses RMTg GABAergic neurons. Furthermore, local activation of trace amine-associated receptor 1 (TAAR1; a trace amine receptor) in the RMTg is sufficient to rescue the despair-like behaviour produced by the loss of AADC expression. Our results identify a novel efferent pathway from the LHb to the RMTg whereby trace aminergic signalling allows the brain to manage acute stress by preventing VTA dopaminergic neuron hypoactivity. The TAAR1-mediated trace aminergic signalling in the LHb-RMTg pathway may hold promise as a therapeutic target for stress-mediated psychological diseases.

## Main

The lateral habenula (LHb) is a behavioural system associated with depression, stress, pain, anxiety, fear, aversive motivation and reward^2–10^. In humans and animals, there is evidence to suggest that this brain region is a powerful negative regulator of dopaminergic systems in the midbrain^1^. The dopaminergic pathway, a major reward-related pathway that projects from the ventral tegmental area (VTA) to the nucleus accumbens (NAc), is important for regulating chronic-stress-induced depression^1,16^.

The current understanding of the neural circuitry between the LHb and the rostromedial tegmental nucleus (RMTg) is that increased activity of LHb glutamatergic neurons might drive augmented RMTg GABAergic neuronal activation, leading to the hypoactivity of VTA dopaminergic neurons^17,18^. However, various types of acute stress excite VTA dopaminergic neurons as well as LHb glutamatergic neurons^11–14^, and little is known about how the activity of the RMTg changes under these circumstances. Since RMTg GABAergic neurons receive a large amount of input from the LHb^19^, acute-stress-induced activation of the LHb and VTA is inconsistent with the known function of the LHb-RMTg-VTA pathway. Therefore, it is important to determine whether the efferent pathway from the LHb to the RMTg plays an inhibitory role as a paradoxical effect of acute stress.

Interestingly, the ‘D-neurons’ in the LHb express L-amino acid decarboxylase (AADC) and produce trace amines rather than monoamines, such as dopamine and serotonin^15,20^. Notably, trace amine-associated receptor 1 (TAAR1)-mediated signalling inhibits the firing frequency of monoaminergic neurons in the midbrain^21,22^. Hence, we reasoned that LHb D-neurons may have a unique suppressive role in the pathway that regulates RMTg GABAergic neurons via trace aminergic signalling under acute stress.

### Acute stress activates AADC-expressing D-neurons in the LHb

D-neurons are located in various parts of the brain, including the LHb (Extended Data Fig. 1a). To clarify the function of the LHb-RMTg-VTA pathway in response to acute stress (Fig. 1a) and gain molecular insights into the function of D-neurons in the LHb, we first performed fluorescence *in situ* hybridization (FISH) analysis in mice by double-labelling AADC in conjunction with a dopaminergic neuron marker (tyrosine hydroxylase, TH), a serotonergic neuron marker (tryptophan hydroxylase 2, TPH2), and a glutamatergic neuron marker (vesicular glutamate transporter 2, VGLUT2). The majority of AADC mRNA was colocalized with VGLUT2 mRNA (Fig. 1b and d) but not with TH or TPH2 mRNA (Fig. 1c, Extended Data Fig. 1), suggesting that a substantial majority of AADC-expressing cells in the LHb are glutamatergic and nonmonoaminergic D-neurons.

**Figure 1:**
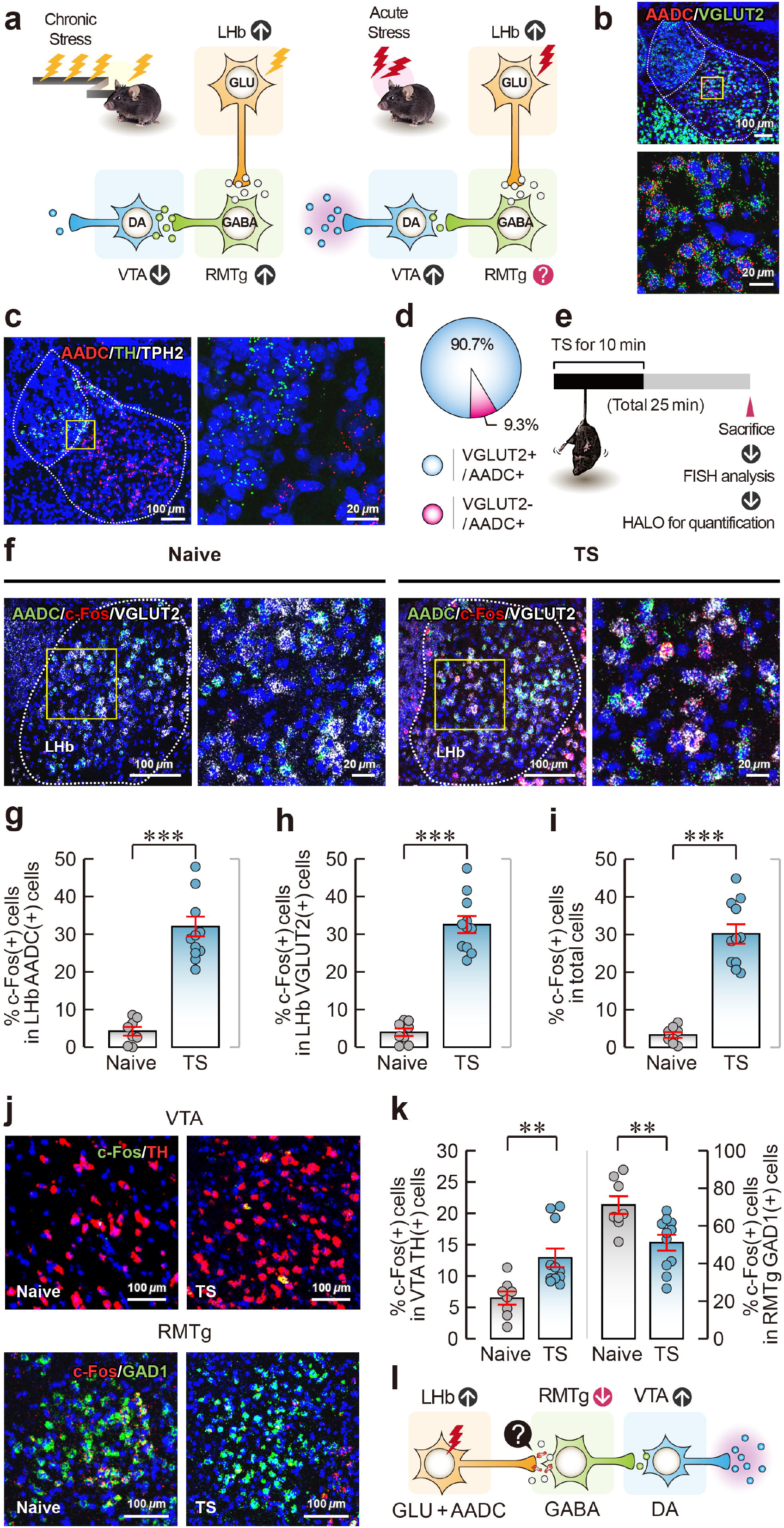
Acute stress alters the neuronal activity of the LHb-RMTg-VTA pathway. **a**, Schematic of changes in neuronal activity of the LHb-RMTg-VTA pathway in chronic and acute stress responses. **b**, **c**, Expression of AADC/VGLUT2 (**b**) and AADC/TH/TPH2 (**c**) in the LHb. **d**, Pie chart of the percentage of VGLUT2-expressing AADC-positive cells. **e**, Experimental schematic of acute tail-suspension-induced stress (TS), which was followed by FISH analysis. **f**, Expression of AADC/c-Fos/VGLUT2 in the LHb of naïve and TS model mice. **g–i**, Percentages of c-Fos-expressing AADC-positive cells (**g**), VGLUT2-positive cells (**h**), and c-Fos-expressing cells in the LHb (**i**). **j**, Expression of c-Fos/TH (top) in the VTA and c-Fos/GAD1 in the RMTg (bottom). **k**, Percentages of c-Fos-expressing TH-positive cells in the VTA (left) and GAD1-positive cells (right) in the RMTg of naïve and TS model mice. **L**, Acute-stress-mediated alterations in neuronal activity in the LHb-RMTg-VTA pathway compared to the activity induced by chronic stress. \**P* < 0.05, \*\**P* < 0.01, and \*\*\**P* < 0.001. Data are presented as the means ±s.e.m. Details on the statistical analyses and sample sizes are provided in Supplementary Table 2.

To evaluate the neuronal activity regulated by acute stress in the LHb-RMTg-VTA pathway, we measured c-Fos expression in the LHb, VTA and RMTg of mice subjected to tail suspension for 10 min. c-Fos expression was increased in glutamatergic neurons, including D-neurons, in the mice exposed to acute stress caused by tail suspension compared to stress-naïve mice (Fig. 1e-I, Extended Data Fig. 2), and no difference in the proportions of c-Fos-expressing glutamatergic neurons and D-neurons was observed (Extended Data Fig. 2). Moreover, acute stress caused by tail suspension increased c-Fos expression in VTA dopaminergic-positive neurons and decreased c-Fos expression in RMTg GAD1-positive neurons (Fig. 1j and k). These findings indicate that acute stress attenuates RMTg GABAergic neuronal activity and augments VTA dopaminergic neuronal activity, despite the increase in LHb neuronal activity; thus, LHb neurons may play an inhibitory role via an as-yet-unknown signalling pathway between the LHb and RMTg (Fig. 1l).

### Activation of LHb D-neurons modulates dopamine secretion through the RMTg-VTA pathway

To investigate the function of LHb D-neurons in acute stress caused by tail suspension, we used a chemogenetic tool. We first confirmed the coexpression of Cre recombinase with endogenous AADC expression in the LHb of *AADC^Cre^* mice (Extended Data Fig. 3a-c). The selective activation of LHb D-neurons in these mice by chemogenetic stimulation significantly lowered the immobility time in the tail-suspension test (TST; Fig. 2a, b) but did not impact locomotion in the open-field test (OFT; Extended Data Fig. 3d and e). Furthermore, c-Fos expression was increased in VTA dopaminergic neurons and decreased in RMTg GABAergic neurons in chemogenetically stimulated mice (Fig. 2c-h, Extended Data Fig. 6c and d). Consistent with the above results, these findings demonstrate that the D-neurons in the LHb regulate the activity of RMTg GABAergic neurons and VTA dopaminergic neurons.

**Figure 2:**
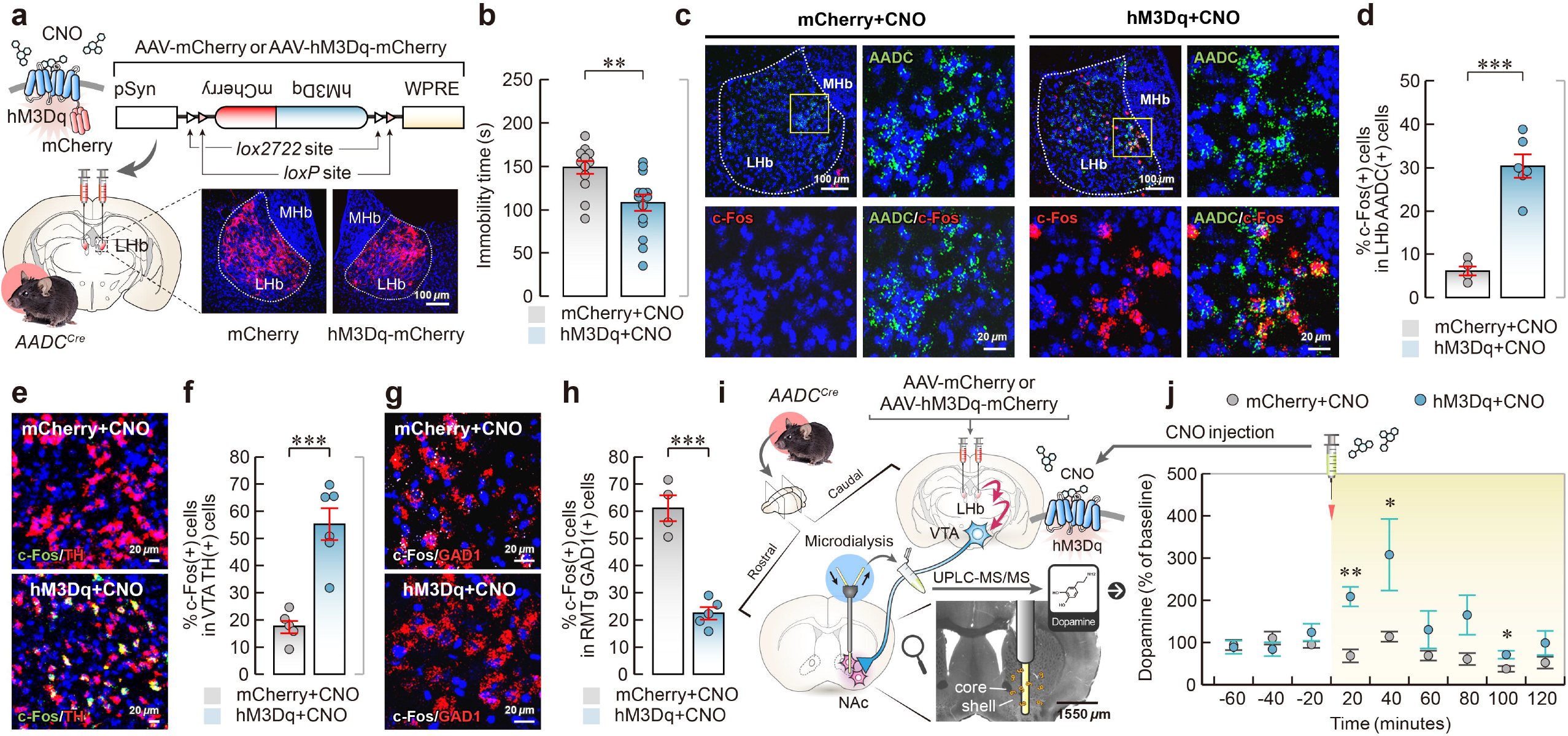
Chemogenetic stimulation of LHb D-neurons regulates dopamine secretion. **a**, Schematic of AAV vectors for Cre-dependent hM3Dq or mCherry expression; illustration of viral injection. **b**, Effect of CNO-induced activation of LHb D-neurons on behaviour observed in the TST. **c–h**, Changes in the expression of c-Fos in VTA dopaminergic neurons and RMTg GABAergic neurons caused by chemogenetic stimulation of LHb D-neurons. Representative FISH images in the LHb (**c**) and the percentage of c-Fos-expressing AADC-positive neurons (**d**). Expression of c-Fos/TH in the VTA (**e**) and c-Fos/GAD1 in the RMTg (**g**). Percentage of c-Fos-expressing TH-positive neurons (**f**) and c-Fos-expressing GAD1-positive neurons (**h**). **i**, **j**, Effect of chemogenetic stimulation of LHb D-neurons on dopamine release in the NAc. Experimental schematic of microdialysis (**i**) and dopamine release changes (**j**). \**P* < 0.05, \*\**P* < 0.01, and \*\*\**P* < 0.001. Data are presented as the means ±s.e.m. Details on the statistical analyses and sample sizes are provided in Supplementary Table 2.

To determine whether the LHb D-neuron activation-induced increase in the activity of VTA dopaminergic neurons affects dopamine secretion, we performed microdialysis in the ventral striatum, including the NAc, of the chemogenetically stimulated mice (Fig. 2i, Extended Data Fig. 3f). Notably, the level of dopamine increased after chemogenetic stimulation of LHb D-neurons (Fig. 2j), which was consistent with the results of c-Fos experiments in VTA dopaminergic neurons (Fig. 2e and f).

### Reduced AADC expression in neurons in the LHb causes depressive-like behaviours

Next, we investigated whether AADC expression is changed in animals with depression induced by chronic restraint stress (CRS) or learned helplessness (LH) protocols using quantitative polymerase chain reaction (qPCR) analysis. The analysis revealed that AADC mRNA was significantly decreased in rats exposed to CRS and in mice exposed to unpredictable electric foot shocks compared to animals that did not undergo stress (Fig. 3a, b). Then, to better understand the role of AADC expression in the LHb in the development of depressive phenotypes, we performed AADC-knockdown experiments in the LHb using adeno-associated virus (AAV)-mediated delivery of a short hairpin RNA (shRNA; shAADC) to reduce AADC mRNA levels (Fig. 3c-e, Extended Data Fig. 4). Depressive phenotypes induced by AADC knockdown were assessed in mice injected bilaterally with shAADC or the control (GFP) virus with behavioural assays (Fig. 3f). Notably, AADC-knockdown mice did not show body weight gain and displayed depressive-like phenotypes, including anhedonia and despair-like behaviours, in the sucrose preference test (SPT) and TST, respectively (Fig. 3g-h). Control and AADC-knockdown mice showed no significant differences in anxiety-like behaviour or locomotion in the OFT or elevated zero maze (EZM; Extended Data Fig. 5).

**Figure 3:**
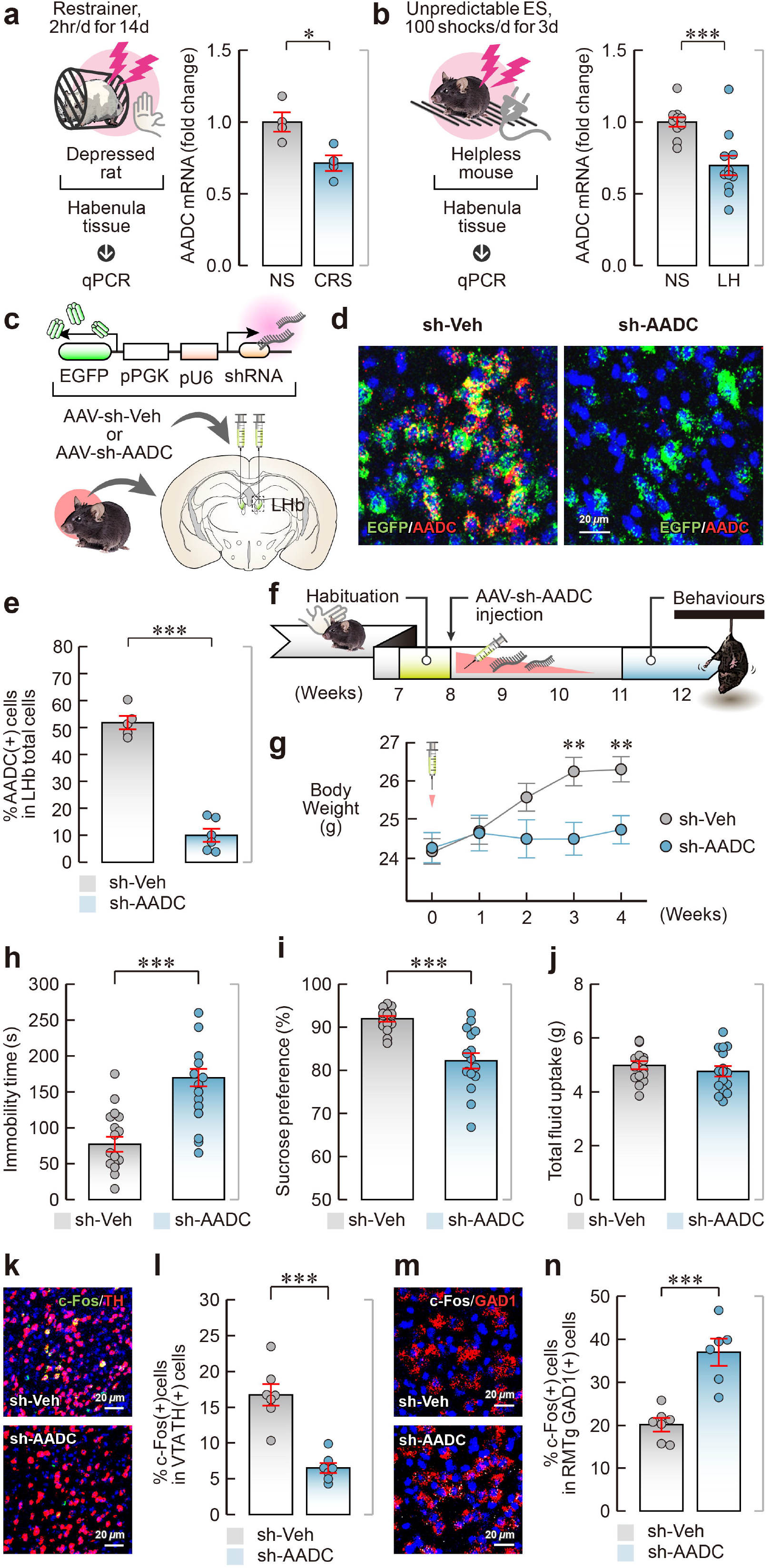
AADC knockdown in the LHb elicits depressive-like behaviours. **a**, **b**, Reduced expression of AADC mRNA in animal models of depression. Experimental schematic (**a**, left, **b**, left), qPCR analysis in rats with CRS (**a**, right) and mice with LH (**b**, right). **c**, Schematic and location of the injection of AAV engineered to overexpress shRNA against AADC. **d**, **e**, *In vivo* validation of AADC knockdown demonstrated by the expression of EGFP/AADC in the LHb (**d**) and the percentage of AADC-positive cells (**e**). **f**, Experimental paradigm for behavioural assays. **g–j**, Effect of AADC knockdown in the LHb on animals’ body weight gain (**g**), immobility time in the TST (**h**), percentage of sucrose preference (**i**), and total fluid intake (**j**) in the SPT. **k–n**, Effect of AADC knockdown in the LHb on VTA dopaminergic and RMTg GABAergic neuronal activity. Expression of c-Fos/TH in the VTA (**k**) and RMTg (**m**). Percentage of c-Fos-expressing TH-positive neurons in the VTA (**l**) and GAD-1-positive neurons in the RMTg (**n**). \**P* < 0.05, \*\**P* < 0.01, and \*\*\**P* < 0.001. Data are presented as the means ±s.e.m. Details on the statistical analyses and sample sizes are provided in Supplementary Table 2.

We next examined c-Fos expression in VTA dopaminergic neurons and RMTg GABAergic neurons in control and AADC-knockdown mice. In contrast to chemogenetic stimulation of LHb D-neurons in mice (Fig. 2e-h), AADC knockdown reduced c-Fos expression in VTA dopaminergic neurons and increased c-Fos expression in RMTg GABAergic neurons (Fig. 3k-n, Extended Data Fig. 6a and b).

### LHb D-neurons form direct synapses with RMTg GABAergic neurons

To identify the neurons innervated by LHb D-neurons, we examined which neurons received direct synaptic input from LHb D-neurons using an anterograde tracing approach (Fig. 4a). Axons of LHb D-neurons labelled with mGFP/synaptophysin-mRuby mostly traversed through the fasciculus retroflexus (fr) and terminated in the RMTg (Fig. 4b and c, Extended Data Movie 1).

**Figure 4:**
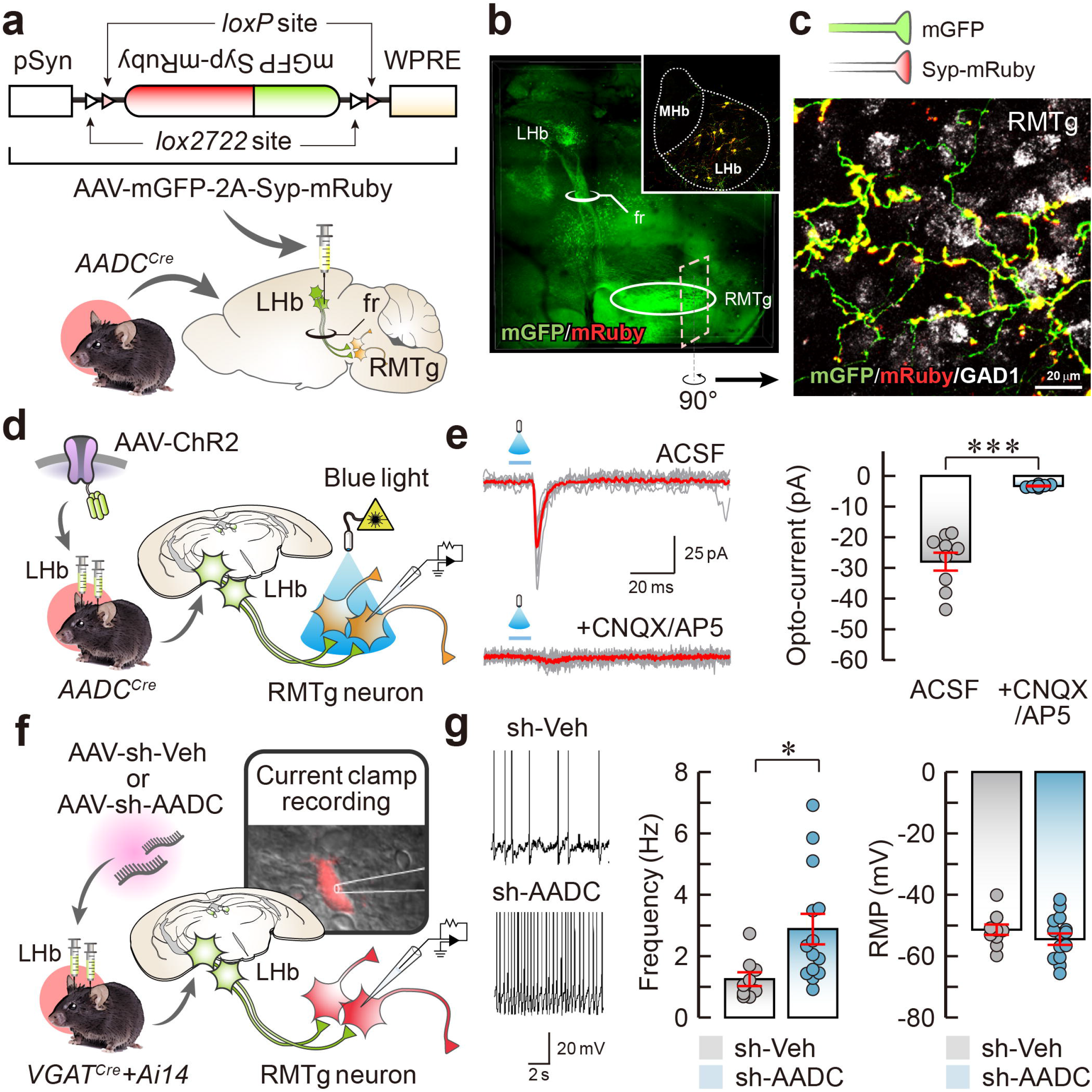
LHb D-neurons innervate RMTg GABAergic neurons. **a**, Schematic of the AAV used and experimental procedure for anterograde tracing. **b**, Representative AAV-hSyn-mGFP-Syp-mRuby infection pattern in the LHb (inset) and sagittal fluorescent image showing LHb D-neuron projections. The LHb, fr, and RMTg are indicated. LHb D-neuronal fibres passing in the fr track and the RMTg. **c**, LHb D-neuronal terminals on GABAergic neurons of the RMTg. **d**, Schematic of AAV-ChR injection into the LHb and the response of RMTg GABAergic neurons to brief optical stimulation. **e**, Traces of oEPSCs (left) and summary data on normalized oEPSC amplitude under baseline conditions after CNQX (20 μM) and AP5 (50 μM) perfusion in ACSF (right). **f**, Schematic of the method for measuring the firing rate of RMTg GABAergic neurons in AADC-knockdown *VGAT^Cre^*::Ai14 mice. **g**, Representative traces (left), firing frequency (middle), and resting membrane potential (right). \**P* < 0.05 and \*\*\**P* < 0.001. Data are presented as the means ±s.e.m. Details on the statistical analyses and sample sizes are provided in Supplementary Table 2.

To further confirm whether LHb D-neurons form direct synaptic contacts, we assessed the activity of RMTg GABAergic neurons using an optogenetic tool and electrophysiology tools (Fig. 4d). Brief optogenetic stimulation of the terminals on LHb D-neurons induced temporally precise inward currents, which were abolished by CNQX and AP5 application (Fig. 4e), suggesting glutamatergic transmission. Thus, we next evaluated whether D-neurons with downregulated AADC expression in the LHb altered the activity of RMTg GABAergic neurons. Electrophysiological recordings of RMTg GABAergic neurons revealed that AADC-knockdown mice had a significantly increased neuronal firing frequency compared with control mice, but the resting membrane potential of neurons in AADC-knockdown mice did not differ from that of control mice (Fig. 4f and g). Therefore, we suggest that glutamatergic LHb D-neurons are inhibitory rather than excitatory to RMTg GABAergic neurons.

### LHb D-neurons regulate RMTg GABAergic neurons by suppressing their activity

To further address whether selective stimulation of LHb D-neurons in chemogenetically stimulated mice suppresses RMTg GABAergic neurons, we measured the firing frequency of RMTg GABAergic neurons using optogenetics. For optogenetic stimulation, a viral vector encoding either shAADC/ChR2 or GFP/ChR2 was bilaterally injected into the LHb of *AADC^Cre^* mice (Fig. 5a). Interestingly, optogenetic stimulation of the nerve terminals of LHb D-neurons in the RMTg significantly reduced the spontaneous firing frequency of RMTg GABAergic neurons in control mice, but this reduction in firing frequency was not observed in AADC-knockdown mice (Fig. 5b, Extended Data Fig. 7a and b). These results indicated that glutamatergic LHb D-neurons have an inhibitory rather than excitatory effect on RMTg GABAergic neurons.

**Figure 5:**
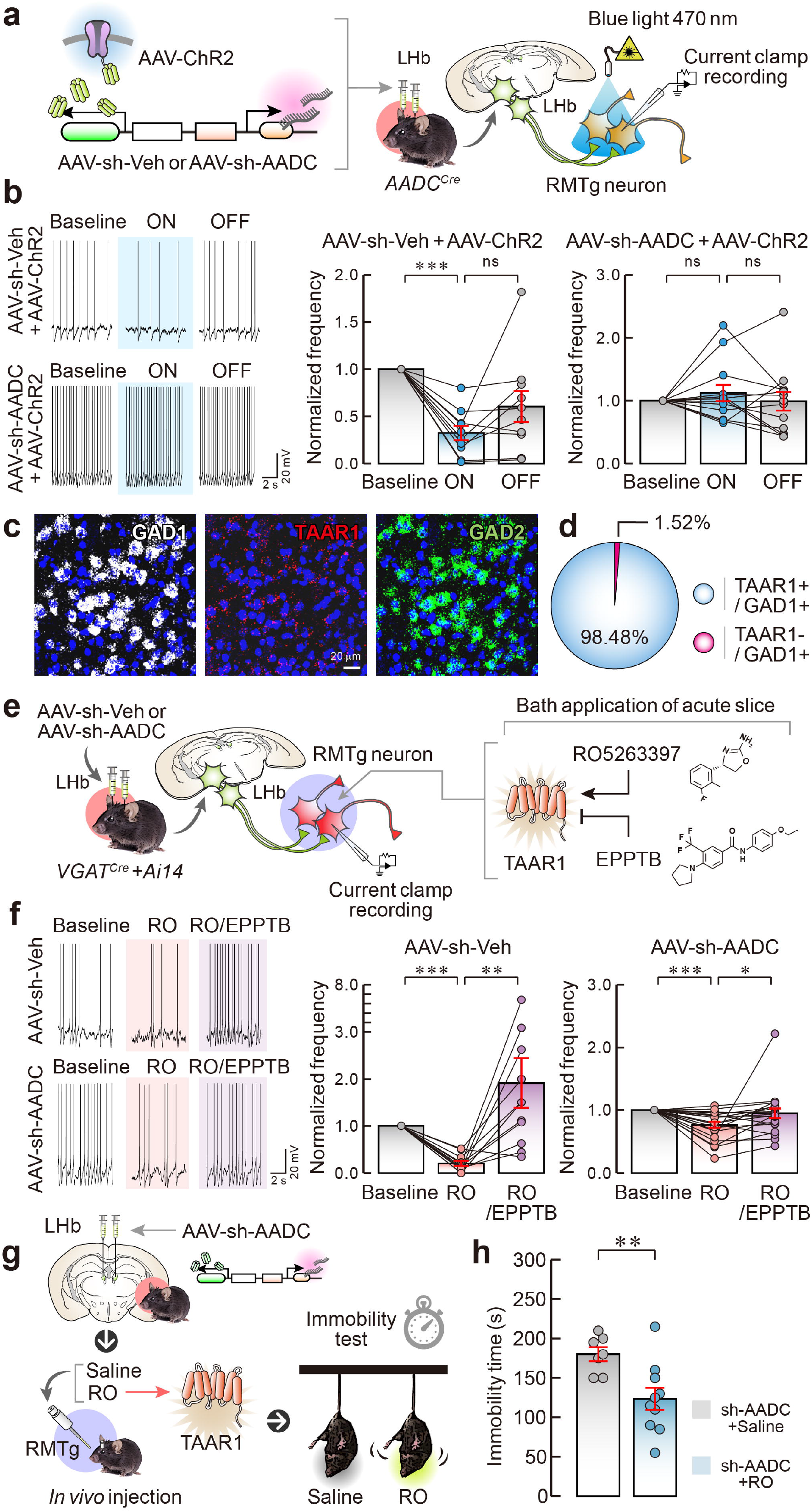
LHb D-neurons suppress RMTg GABAergic neurons through trace aminergic signalling. **a**, Schematic of the AAV used and RMTg GABAergic neuronal activity induced by the photostimulation of LHb D-neurons. **b**, Representative traces (left) and firing frequency in the control mice (middle) and AADC-knockdown mice (right) measured in the ON or OFF phases of blue light photoactivation. **c**, **d**, Expression of TAAR1/GAD1 mRNA in GABAergic neurons of the RMTg (**c**) and the percentage of TAAR1-expressing GAD1-positive GABAergic neurons in the RMTg (**d**). **e**, Schematic of AAV injection and RMTg GABAergic neuronal firing rates in response to the application of RO5263397 or EPPTB. **f**, Representative traces (left) and firing frequency in control (middle) and AADC-knockdown mice (right) measured in response to the application of RO5263397 (500 nM) or EPPTB (1 μM). The firing frequency was normalized to the baseline value (**b**, **f**). **g**, **h**, Effect of RO5263397 application in AADC-knockdown mice *in vivo* on their performance in the TST. Schematic of an AAV *in vivo* injection (**g**) and TST immobility time results (**h**). \**P* < 0.05, \*\**P* < 0.01, and \*\*\**P* < 0.001. Data are presented as the means ±s.e.m. Details on the statistical analyses and sample sizes are provided in Supplementary Table 2.

Given that the selective activation of D-neurons in the LHb plays an inhibitory role, we next explored the molecular pathway whereby D-neurons mediate the suppression of RMTg GABAergic neurons. Considering that D-neurons produce trace amines but no monoamines, trace aminergic signalling may play a major role in LHb D-neurons in the RMTg GABAergic neuronal circuit. TAAR1 is the main receptor for trace amines in the brain^23^. Thus, to determine whether RMTg GABAergic neurons express TAAR1, we examined these neurons using FISH. Approximately 98% of GAD1-positive GABAergic neurons in the RMTg expressed TAAR1 mRNA (Fig. 5c and d, Extended Data Fig. 8), which suggests that most RMTg GABAergic neurons are affected by trace amines.

Next, to assess the effect of trace amines on RMTg GABAergic neurons, slices containing the RMTg from control and AADC-knockdown mice were acutely exposed to a selective TAAR1 agonist ((4S)-4-(3-fluoro-2-methylphenyl)-4,5-dihydro-2-oxazolamine hydrochloride, or RO5263397) or antagonist (N-(3-ethoxy-phenyl)-4-pyrrolidin-1-yl-3-trifluromethyl-benzamide, or EPPTB) by bath application (Fig. 5e). Bath application of RO5263397 (500 nM) suppressed the spontaneous firing frequency of RMTg GABAergic neurons in slices from control mice. In contrast, additional application of EPPTB (1 μM) not only inhibited the RO5263397-mediated decrease in firing frequency but also significantly increased the firing frequency of these neurons above the basal level (Fig. 5f, Extended Data Fig. 7c). In AADC-knockdown mice, RO5263397 also significantly decreased the firing frequency of RMTg GABAergic neurons, but additional application of EPPTB restored the firing frequency of these neurons to the basal level (Fig. 5f, Extended Data Fig. 7d). Together, these findings indicate that trace aminergic signalling between glutamatergic LHb D-neurons and RMTg GABAergic neurons acts as a powerful inhibitor.

To determine the effect of trace amines on RMTg GABAergic neurons *in vivo*, we administered RO5263397 or saline directly into the RMTg in AADC-knockdown mice and subjected them to the TST (Fig. 5g). The immobility time in the TST was rescued in RO5263397-treated AADC-knockdown mice compared with saline-treated knockdown mice (Fig. 5h). This result shows that *in vivo* administration of the TAAR1 agonist led to a full recovery of the despair-like behaviour induced by the lack of trace amines induced by AADC knockdown.

### Conclusion

LHb neuron hyperexcitability suppression has emerged as a major target for the restoration of the LHb-midbrain monoamine pathway to its normal condition. As part of this effort, several researchers have attempted to elucidate the existence and role of GABAergic neurons within the LHb^24–28^. However, the presence and function of GABAergic neurons capable of inhibiting LHb hyperexcitability were insufficient. Even though LHb D-neurons are glutamatergic neurons, we revealed that these cells can sufficiently perform inhibitory functions using trace amines in the well-known LHb-RMTg-VTA pathway and that this inhibitory role plays a defence function under acute stress. Trace aminergic output from LHb D-neurons suppresses RMTg GABAergic neurons; such suppression is strengthened by acute stress under normal conditions to increase the activity of VTA dopaminergic neurons, thereby preventing rapid onset of depressive phenotypes (Extended Data Fig. 9).

We classified LHb glutamatergic neurons into pure glutamatergic neurons and D-neurons. Optogenetic stimulation of distinct pathways projecting from the medial prefrontal cortex^29^ and lateral hypothalamus^13,30,31^ to LHb glutamatergic neurons showed the opposite effect on escape-related behaviour. These contradictory results need further investigation to close the gap between studies exploring behavioural paradigms of the LHb. The inhibitory role of LHb D-neurons can explain this discrepancy and present a new paradigm for understanding various functions of the LHb.

Trace aminergic signalling has attracted attention as a new therapeutic target due to the reduction in trace amines in psychiatric disorders, including depression and schizophrenia^32–37^. Because trace amines have the limitations of short half-lives and unusual characteristics for storage, transport, and diffusion^38^, many researchers focus on molecular function studies on TAARs to understand trace aminergic signalling^22,39,40^. Our findings overcome these limitations and provide new insight that could help clarify the as-yet-unknown role of the remaining D-neurons in the brain. Further studies on the neural circuits between numerous D-neurons and TAAR-expressing neurons in the brain might promote an understanding of the brain functions involved in trace aminergic signalling.

## Acknowledgements

We thank NeuroVis Inc. for performing the *in vivo* measurement of dopamine release, and Ulsan National Institute of Science and Technology Optical Biomed Imaging Center (UOBC) for three-dimensional analysis of the neural circuit. This work was supported by the National Research Foundation of Korea (NRF), funded by the Ministry of Science, ICT & Future Planning (NRF-2017M3C7A1079692 to H.K.; NRF-2017R1D1A1B06032730 to H.W.L.; and E0210201-01 to J.L.). The authors declare that they have no conflicts of interest.

## Author contributions

S.H.Y., H.W.L. and H.K. designed the study. E.Y. performed the FISH analysis and histology experiments. J.L. and S.C. performed the *in vitro* patch-clamp experiments. H.Y. and H.S.P. produced the viruses and carried out the qPCR analysis. S.H.Y. performed the stereotaxic surgery. S.H.Y., J.Y.K. and J.T.J. conducted the behavioural experiments and established the animal stress models. D.L. analysed the experimental data and drew the schematics. H.W.L. and H.K. supervised all aspects of the work. S.H.Y., H.W.L. and H.K. wrote the manuscript with input from all authors.

## Methods

### Animals

Adult male C57BL/6J mice and Sprague–Dawley rats (7 weeks of age) were purchased from Japan SLC Inc. (Shizuoka) and were used after habituation for one week. *AADC^Cre^*(B6.FVB(Cg)-Tg(Ddc-cre)SD56Gsat/Mmucd, RRID:MMRRC_037410-UCD) mice were obtained from the Mutant Mouse Resource and Research Center (MMRRC) of the University of California-Davis (Davis) and were used for chemogenetics, optogenetics, microdialysis, and anterograde tracing experiments. *VGAT^Cre^* (B6J.129S6(FVB)-Slc32a1^*tm2(cre)LoW1*^/MwarJ, RRID:IMSR_JAX:028862)^41^ mice were crossed with Ai14 (B6. Cg-Gt(ROSA)26Sor^*tm14(CAG-tdTomato)Hze*^/J,RRID:IMSR_JAX:007914)^42^ mice to identify GABAergic neurons in the RMTg. All animals were housed three to four per cage under a 12/12-h light/dark cycle (lights on at 8 a.m.) and given *ad libitum* access to food and water. Animals were habituated to the facility for at least 1 week prior to the experiments. All animal experiments were approved by the Korea University Institutional Animal Care and Use Committee (IACUC) and performed in accordance with the guidelines of Korea University (Study approval number KOREA-2017-0007-C1).

### FISH analysis

*In situ* hybridization was performed according to the procedure described by Yang *et al.*^43^. In brief, frozen sections (14 μm thick) were cut coronally through the Hb, VTA, and RMTg. Sections were thaw-mounted onto Superfrost Plus Slides (#12-550-15, Fisher Scientific). Then, the sections were fixed in 4% paraformaldehyde (PFA) for 10 min, dehydrated in increasing concentrations of ethanol for 5 min, and finally air-dried. Tissues were then pretreated for protease digestion for 10 min at room temperature. Probe hybridization and amplification were performed at 40 °C using a HybEZ hybridization oven (Advanced Cell Diagnostics). The probes used in this study are described in Supplementary Information Table 1. The labelled probes were conjugated to Alexa Fluor 488, Atto 550, and Atto 647. The sections were hybridized with the labelled probe mixture at 40 °C for 2 h. Unbound hybridization probes were removed by washing the sections three times with 1× wash buffer at room temperature for 2 min; then, the slides were treated with Amplifier 1-FL for 30 min, Amplifier 2-FL for 15 min, Amplifier 3-FL for 30 min, and Amplifier 4 Alt B-FL for 15 min. After each amplifier solution treatment session, sections were washed with 1× wash buffer at room temperature for 2 min before being treated with the next amplifier solution. Next, the slides were viewed, analysed, and photographed using a TCS SP8 Dichroic/CS microscope (Leica). After FISH was performed, the average number of dots per cell was quantified using the HALO image analysis algorithm in HALO v2.3.2089.18 (Indica Labs) software^44^.

### Histology

Mice were anaesthetized and perfused with 0.9% saline, followed by perfusion with 4% PFA. The brains were removed from the skulls and then postfixed in 4% PFA overnight at 4°C. After postfixation, the brains were incubated in 30% sucrose at 4°C. With a cryotome (Leica CM300, Leica), the brains were cut into 100 μm thick coronal sections; sectioned brain regions encompassed the habenula. Next, sections were washed with phosphate-buffered saline (PBS) three times for 5 min at room temperature. The washed sections were incubated with Hoechst (H3570, Invitrogen), a blue fluorescent stain that labels DNA, at room temperature for 10 min.

The stained sections were immersed in mounting solution for 30 min at 37 °C. The sections were viewed and photographed using a TCS SP8 Dichroic/CS microscope (Leica).

### Stress models

Acute stress was induced in mice by tail suspension as previously described^14^. After 1 week of habituation to their home cages, experimental mice were suspended by the tail for 10 min. The mice were sacrificed 25 min after the onset of tail suspension, and samples were collected for FISH analysis.

CRS was induced as previously described^45^. In brief, eight-week-old Sprague–Dawley rats were subjected to restraint stress in DecapiCones^®^ rodent restrainers (MSPP-DCL120, Braintree Scientific) for 2 h per day for 2 weeks. Control rats (no stress, NS) were maintained in their home cages without being disturbed.

LH was assessed as previously described^46^. Briefly, eight-week-old C57BL/6J mice were placed in shock chambers (chamber dimensions, 30 cm wide × 30 cm deep × 25 cm high; Multi Conditioning System, TSE system), where they underwent 100 inescapable electric foot shocks of 0.3 mA intensity and 5 sec duration with intershock intervals of 5-99 sec.

### qPCR

qPCR was performed using the same protocol as a previous study^45^. Briefly, the habenula was isolated from rats (10 weeks of age) and mice (8 weeks of age), and the RNA from the habenula tissue was subjected to reverse transcription. The reverse-transcribed sequences were then amplified by qPCR, and the products were electrophoresed using 2%agarose gels. The comparative Ct method (ΔΔCt) was used for the relative quantification of the amplification products and to calculate the fold changes in gene expression between rats that underwent CRS and those with NS^47^ and between naïve mice and those that completed the LH experiment. The expression levels of AADC were normalized to the expression level of the housekeeping gene GAPDH. The sequences for the specific primers were as follows: for the CRS rat model, AADC forward, 5’-TTCTTCGCTTACTTCCCCACG-3’; AADC reverse, 5’-CCCAGCCAATCCATCATCACT-3’; GAPDH forward, 5’-CATCCACTGGTGCTGCCAAGGCTG-3’; and GAPDH reverse, 5’-ACAACCTGGTCCTCAGTGTAFCCCA-3’ and for the LH mouse model, AADC forward, 5’-GGCTTACATCCGAAAGCACG-3’; AADC reverse, 5’-CTTTAGCCGGAAGCAGACCA-3’; GAPDH forward, 5’-ACCCAGAAGACTGTGGATGG-3’; and GAPDH reverse, 5’-CACATTGGGGGTAGGAACAC-3’.

### Viruses

The virus production methodology used in this study to achieve AADC knockdown was based on a previously described protocol^45^ but was modified such that recombinant AAV serotype 2/9 was used, and the virus titres were determined by real-time PCR. AAV plasmids expressing hM3Dq [pAAV-hSyn-DIO-hM3D(Gq)-mCherry (#44361)], mCherry [pAAV-hSyn-DIO-mCherry (#50459)], mGFP/synaptophysin-mRuby [pAAV-hSyn-FLEx-mGFP-2A-Synaptophysin-mRuby (#71760)] and channel rhodopsin (ChR) [pAAV-EF1a-double floxed-hChR2(H134R)-EYFP-WPRE-HGHpA (#20298)] were purchased from Addgene. The viral vector was constructed using GFP [pAAV-U6-GFP (provided by Cell Biolabs)] with the shRNA sequence against *AADC* mRNA (shAADC, 5’-GTGATCTAGCAAGCAGTGT-3’) inserted to knock down AADC expression. A virus encoding only GFP and with no shRNA was used as a control. For AADC-knockdown validation, full-length complementary DNA (cDNA) of AADC was transcribed from a C57BL/6N mouse brain cDNA library using RT–PCR and inserted into a pDEST-GFP plasmid. For transfection, HEK293T cells were plated on 60 mm dishes and cultured for 24 h. Plasmids encoding GFP-AADC, AADC, and scrambled shRNAs were transfected into the cells using Lipofectamine (GIBCO). Seventy-two hours after transfection, HEK293T cells were suspended in RIPA lysis buffer (T&I) containing a protease inhibitor cocktail (Roche). After centrifugation, cell lysates were cleared of cell debris, and the remaining components were separated by 10% sodium dodecyl sulfate–polyacrylamide gel electrophoresis (SDS–PAGE) before being transferred to polyvinylidene fluoride (PVDF) membranes. Membranes were immunoblotted overnight at 4°C with anti-GFP (1:1000; B-2, Santa Cruz) as the primary antibody. The membranes were washed, incubated with species-specific horseradish peroxidase–conjugated secondary antibody for 1 h at room temperature and developed using electrochemiluminescence (ECL) solution (Thermo Scientific Pierce).

### Stereotaxic surgery

Male adult mice (8 weeks of age) were anaesthetized with isoflurane (5% induction, 1% maintenance) and placed on a stereotaxic apparatus (Ultra Precise stereotaxic instruments for mice; Stoelting Co.). After an incision was made in the scalp, a craniotomy was performed using a hand drill so that the virus designed to achieve AADC knockdown could be injected into the LHb. Approximately 1 μL of the virus was injected into the LHb (coordinates from bregma: −1.58 mm anterior/posterior (A/P), ±0.9 mm medial/lateral (M/L), −3.1 mm dorsal/ventral (D/V), 10° angle towards the midline in the coronal plane) using a microinjection cannula (30 gauge, Plastics One) and Ultra Micro Pump III (World Precision Instruments) at a speed of 0.12 μL/min. The incision was closed with 9 mm autoclips (#205016, MikRon Precision Inc.), and antibiotics and analgesics were administered to the mice. Mice were placed in a clean cage on a heating pad and allowed to recover from anaesthesia. Then, they were kept in their home cage for 3 weeks so that the AAV could take effect before behavioural tests were performed.

For chemogenetic stimulation experiments, recombinant AAVs expressing hM3Dq or mCherry were bilaterally injected into the LHb of *AADC^Cre^* mice. A CMA7 guide cannula (CMA microdialysis) was implanted to measure dopamine in the ventral striatum (+1.00 mm A/P, +1.60 mm M/L and −3.00 mm D/V), including the NAc. Mice were injected with clozapine N-oxide (CNO; BML-NS105, Enzo), prepared in sterile 1× PBS with 0.5% dimethyl sulfoxide (DMSO; SHBD9284V, Sigma–Aldrich), at 5 mg/kg of body weight for hM3Dq 40 min before the start of behavioural testing.

For knockdown experiments, recombinant AAVs expressing GFP or shAADC were bilaterally injected into the LHb of wild-type mice. A guide cannula (26 gauge, Plastics One) was implanted to enable the administration of a TAAR1 agonist in the RMTg (−4.24 mm A/P, +0.70 mm M/L, −4.00 mm D/V and 10° angle), and mice were allowed to recover for 1 week after implantation.

For anterograde tracing experiments, recombinant AAVs expressing mGFP/synaptophysin-mRuby were bilaterally injected into the LHb of *AADC^Cre^* mice. After 4 weeks, mice were sacrificed for histology.

For optogenetic stimulation experiments, recombinant AAVs expressing ChR, ChR with GFP or ChR with shAADC were bilaterally injected into the LHb of *AADC^Cre^* mice. After 3 weeks, the mice were sacrificed to obtain tissues for electrophysiology.

For pharmacological experiments, recombinant AAVs expressing GFP or shAADC were bilaterally injected into the LHb of *VGAT^Cre^*::Ai14 mice. After 3 weeks, the mice were sacrificed to obtain tissues for electrophysiology.

### Behavioural assays

All behavioural assays were performed during the light phase except for the SPT, which was performed during the dark phase to maximize the consumption of solution. In all behavioural experiments, the experimenter was blinded to the animals’ genotypes and the experimental conditions; the data were analysed in a blinded manner as well.

#### OFT

Each mouse was placed in the corner of the open-field chamber (45 cm × 45 cm × 40 cm) and allowed to explore for 15 min. The total distance moved, time spent in the centre, frequency of visits to the centre, and latency to visit the centre were all calculated using EthoVision XT 12 tracking software (Noldus).

#### EZM

Mice were singly placed in a closed quadrant and allowed to explore the apparatus freely for 5 min. The apparatus consisted of two open quadrants and two closed quadrants and was elevated 60 cm above the floor. The total distance travelled, time spent in the open quadrants, frequency of visits to the closed quadrants, and latency to visit the open quadrants were all recorded by EthoVision XT 12 tracking software.

#### TST

The TST was conducted using a 4-chamber apparatus divided by opaque, matte-surfaced acrylic partitions. Each mouse was suspended by their tail in one of the chambers using adhesive tape. A video was recorded for 6 min, and the last 4 min was scored for the immobility time. Immobility was determined every 5 sec, and the immobility time was calculated by (number of instances of immobility × 5 sec). Any instance when the mouse did not move any of its limbs was scored as immobility.

#### SPT

The SPT was conducted using a previously described procedure^48^ with modifications. Singly housed mice were habituated to two water bottles containing tap water for a day and then presented with two identical bottles, one with 1% sucrose solution and the other with tap water. To minimize the potential effect of side preference, the positions of the two bottles were switched daily. Sucrose and water consumption were recorded daily by reweighting the preweighed bottles of test solutions. Sucrose preference was calculated as a relative ratio (mass of sucrose solution intake/total fluid intake).

### Microdialysis

To examine the extracellular level of dopamine *in vivo*, a CMA7 microdialysis probe with a 2 mm membrane length (#P000083, CMA Microdialysis) was used. The probe was slowly inserted into the ventral striatum of mice through guide cannulas and connected to a single-channel liquid swivel and a counterbalancing system 2 h before the experiments. The probe was perfused with sterile artificial cerebrospinal fluid (ACSF) from CMA Microdialysis (147 mM NaCl, 2.7 mM KCl, 1.2 mM CaCl_2_, 0.85 mM MgCl_2_) at a flow rate of 1.0 μl/min with a microinfusion pump. The perfusate was collected every 20 min with a refrigerated microfraction collector; the first 3 times, perfusate was collected for baseline measurements after 2 h of preperfusion, and then perfusate was collected 6 times after the injection of CNO 5 mg/kg (0.5% DMSO, 1× PBS). Dopamine quantification was performed using an ultrahigh-performance liquid chromatography–tandem mass spectrometry (UPLC–MS/MS) system. This system consisted of a SCIEX ExionLC system with a Waters Acquity HSS T3 column (2.1 × 100 mm, 1.8 μm) and a 6500+quadrupole ion trap (QTRAP) mass spectrometer with an electrospray ionization (ESI) source. The data were acquired and quantified using Analyst software version 1.7. The mobile phase consisted of 5 mM ammonium formate and 0.1% formic acid in water and 5 mM ammonium formate in acetonitrile:methanol (v/v, 1:1). The flow rate was 0.3 mL/min, and the injection volume was 10 μL. The mass spectrometer was optimized, and a multiple reaction monitoring (MRM) scan was performed in positive ion mode.

### Circuit tracing

To track mGFP/synaptophysin-mRuby neurons or terminals, mice were perfused with 0.9% saline, followed by 4% PFA at 4 weeks after virus injection. The brains of the mice were removed from the skull and then postfixed in 4% PFA overnight at 4 °C. Fixed brains were cut into 2 mm thick midline sagittal sections, including the LHb and the RMTg, using a sagittal brain matrix for mice (Harvard Apparatus). Sagittal brain sections were optically cleared by a Tissue Clearing Kit (HRTC-001 and SHMS-060, Binaree). Briefly, brain sections were incubated in starting solution at 4 °C and then washed with distilled water three times, 1 h each time, at 4 °C. The washed brain sections were incubated with the tissue-clearing solution in a shaking incubator at 37 °C for 4 days and then incubated in mounting and storage solution. A z-stack of the sections was acquired on a z.1 light sheet fluorescence microscope (Carl Zeiss), and the images were three-dimensionally reconstructed using Imaris software (Bitplane).

For the characterization of presynaptic neurons in the LHb and postsynaptic neurons in the RMTg, FISH was performed using free-floating brain sections that included the LHb and the RMTg. For this analysis, the perfused and postfixed brains were equilibrated in RNase-free 30% sucrose and then cut into 40-μm-thick coronal sections on a cryotome (Leica CM300, Leica). Floating brain sections were placed in chamber slides (Thermo Fisher Scientific) containing pretreatment 2 (ACD) preheated to 60 °C and were then incubated for 10 min to allow sucrose cross-link digestion. The brain sections were then pretreated with pretreatment 4 (ACD) to allow protease digestion at room temperature for 30 min. Probe (GAD1) hybridization, amplification, and analysis were conducted as described above (FISH).

### Electrophysiology

For slice preparation, mice were euthanized by isoflurane inhalation, and using an oscillating tissue slicer (Leica VT1000s, Leica), 300 μm thick coronal brain slices containing the RMTg area were obtained in cold cutting solution with the following composition (in mM): 92 N-methyl-D-glucamine (NMDG), 2.5 KCl, 1.25 NaH_2_PO_4_, 30 NaHCO_3_, 30 HEPES, 0.5 CaCl_2_, 10 MgCl_2_, and 25 glucose, aerated with 95% O_2_/5% CO_2_. The slices were allowed to recover at room temperature for at least 1 h in a recovery solution with the following composition (in mM): 92 NaCl, 2.5 KCl, 1.25 NaH_2_PO_4_, 30 NaHCO_3_, 30 HEPES, 2 CaCl_2_, 2 MgCl_2_, and 25 glucose, saturated with 95% O_2_/5% CO_2_. Each slice was transferred from a recovery reservoir to the recording chamber of a fixed-stage upright microscope (Olympus BX51WI, Olympus) and perfused with oxygenated ACSF with the following composition (in mM): 124 NaCl, 2.5 KCl, 1.25 NaH_2_PO_4_, 24 NaHCO_3_, 1.5 CaCl_2_, 1.5 MgCl_2_, and 10 glucose, saturated with 95% O_2_/5% CO_2_. ACSF was supplied to the chamber at a rate of 1.5–2 mL/min. Each submerged slice was visualized either directly via the microscope’s optics or indirectly via a high-resolution charge-coupled device (CCD) camera system (optiMOS, Qimaging) receiving output from a CCD camera attached to the microscope’s video port.

Whole-cell patch-clamp recordings in current-clamp mode were used to measure spontaneous action potential (AP) firing in RMTg GABAergic neurons indicated by red fluorescence in *VGAZ^Cre^*::Ai14 mice. The recordings were obtained using borosilicate glass pipettes (resistance 4–10 MΩ) prepared using a 2-stage vertical pipette puller (P-1000, Sutter Instrument, or PP-83, Narishige). The pipettes were filled with a solution containing the following (in mM): 140.0 K-gluconate, 10.0 HEPES, 0.5 EGTA, 10.0 glucose, 2.0 Na-ATP, and 0.5 Na-GTP; the pH was adjusted to 7.2 with KOH. Signals were amplified and filtered (2 kHz) using a MultiClamp 700B amplifier, sampled at 5 kHz using Digidata 1440, and recorded using pClamp10 software (Molecular Devices, Union City, CA, USA). No correction for liquid junction potential was made. Analyses were performed using Clampfit 10 (Molecular Devices). Only cells with stable access resistance of < 25 MΩ were included in the analysis.

To examine whether the RMTg directly receives synaptic input from LHb D-neurons, we investigated the responses of RMTg neurons using optogenetic stimulation in *AADC^Cre^* mice. For brief stimulation (10 ms), blue light (473 nm, 0.5 ~ 1 sec, and ~ 10 mW) was delivered by a light-emitting diode (LED) light source (PSU-III-LED, Optoelectronics Tech. Co. Ltd) via an optic fibre during whole-cell recordings from RMTg neurons. Recording pipettes were filled with a solution containing the following (in mM): 127 CsMeSO_4_, 10 NaCl, 5 EGTA, 4 Mg-ATP, 2 Na-GTP, and 2 QX314. To confirm glutamatergic inputs, optically evoked excitatory postsynaptic currents (oEPSCs) were recorded during the application of the competitive AMPA/kainate receptor antagonist 6-cyano-7-nitroquinoxaline-2,3-dione (CNQX; 20 μM, Tocris) and the selective NMDA receptor antagonist amino-5-phosphonopentanoic acid (AP5; 50 μM, Tocris).

For optogenetic stimulation experiments to confirm the function of LHb D-neurons using *AADC^Cre^* mice, blue light (473 nm, 0.5 ~ 1 sec, and ~ 10 mW) was delivered by an LED light source via an optic fibre during whole-cell recordings from RMTg neurons. Blue light was applied between 60 ~ 80 sec and then turned off.

For pharmacological experiments on RMTg GABAergic neurons using control mice and AADC-knockdown mice, APs were recorded under the effects of the TAAR1 agonist RO5263397 (500 nM, Tocris) and the TAAR1 antagonist EPPTB (1 μM, Tocris) in ACSF. All drugs were dissolved in 1% DMSO and stored at −70 °C before use. Recordings began at least 2 min after whole-cell recording was established via break-in. The frequency of AP firing was analysed based on a 30-sec period after drug application.

### Drug administration

For *in vivo* administration, RO5263397 was dissolved in sterilized 1× PBS at a concentration of 1 μg/μL. The drug or vehicle was injected into the RMTg of AADC-knockdown mice through the cannula using an Ultra Micro Pump III at a speed of 0.3 μL/min. Thirty minutes after drug or vehicle injection, the mice were subjected to the TST for 6 min.

### Statistical analysis

Samples were excluded from analyses if the viral injection sites were outside the LHb. Statistical analysis was conducted with IBM SPSS Statistics 25 for Windows (IBM). Pairwise comparisons between 2 groups were performed using a two-tailed independent Student’s *t* test or the Mann– Whitney U test. For comparisons among more than 2 groups, analysis of variance (ANOVA) was used with the appropriate *post hoc* tests. For body weight data, repeated-measures ANOVA was conducted with *post hoc* tests. To monitor changes over time, repeated-measures ANOVA was run, followed by a contrast test. Data are expressed as the mean ± s.e.m., and *P* < 0.05 was considered significant. Details of the statistical analysis, including the numbers of animals, the exact statistical tests used, and the analysis results, are reported in the Source Data.

## Data availability

The data that support the findings of this study are available from the corresponding author upon reasonable request.

## Extended data figure legends

**Extended Data Figure 1: Validation of FISH probes and the characterization of LHb D-neurons a**, Distribution of D-neurons in the brain^49^. D1, spinal cord; D2, nucleus tractus solitarius; D3, parabrachial complex (rostral medulla and pons); D4, midbrain (nuclei associated with the posterior commissure); D5, pretectal nuclei; D6, lateral habenula; D7, paracentral nucleus of the dorsal thalamus; D8, nucleus premammillaris of the hypothalamus; D9, arcuate nucleus; D10, zona incerta; D11, lateral hypothalamic region; D12, dorsomedial hypothalamic nucleus; D13, suprachiasmatic nucleus (SCN); D14, bed nucleus of the stria terminalis; D15, striatum; D16, nucleus accumbens; D17, basal forebrain; D18, cerebral cortex. **b–d,**Expression of VGLUT1/VGLUT2 in the habenula (Hb). VGLUT2 mRNA in both the medial habenula (MHb) and the LHb and (**c**) VGLUT1 mRNA only in the MHb (**d**). **e–h**,AADC mRNA also colocalized with TPH2-positive serotonergic neurons in the raphe nucleus. Neither TH nor TPH2 mRNA was expressed in AADC-expressing D-neurons in the LHb, and 66.32% of VGLUT2-positive neurons were AADC-positive (**i**, **j**). Details on the sample sizes are provided in Supplementary Table 2.

**Extended Data Figure 2: FISH analysis of activated neurons in the LHb after exposure to tail-suspension stress. a**, Percentage of AADC-expressing cells among c-Fos-positive cells. **b**, Percentage of VGLUT2-expressing c-Fos-positive cells. **c**, Average total number of c-Fos copies per μm^2^ in the LHb. \**P* < 0.05. TS, tail suspension. Data are presented as the means ±s.e.m. Details on the statistical analyses and sample sizes are provided in Supplementary Table 2.

**Extended Data Figure 3: Validation of *AADC^Cre^* transgenic mice, effects of chemogenetic stimulation of LHb D-neurons on locomotor activity, and the localization of microdialysis probes. a–c**, Schematic for the validation of *AADCC^re^* mice using the Cre-dependent virus (a recombinant AAV expressing mGFP/synaptophysin-mRuby) **(a)**. Representative image of mGFP and AADC mRNA expression in the LHb **(b)**. Pie chart depicting the proportion of AADC-expressing cells among mGFP-positive cells **(c)**. **d**, **e**, Travel distance (**d**) and velocity (**e**) of mice in the OFT after chemogenetic stimulation of LHb D-neurons. No significant difference was detected. **f**, Schematic drawing of the microdialysis probe placement in the NAc used in Figure 2**i**, **j**. Details on statistical analyses and sample sizes are provided in Supplementary Table 2.

**Extended Data Figure 4: Validation of AADC knockdown in the LHb. a**, **b**, *In vitro* validation of AADC knockdown in HEK293T cells expressing GFP-AADC (pDEST-GFP-AADC) and shVeh or AAV-shAADC plasmid. After 72 h of expression, Western blot analysis (**a**) and the band intensity of exogenous AADC protein levels normalized to that of actin (**b**). **c**, **d**, *In vivo* validation of AADC knockdown in LHb cells expressing AAV-shVeh or AAV-shAADC for 3 weeks. Representative *in situ* hybridization images of EGFP and AADC (**c**). Magnified images of the region of interest are shown in Figure 3**e**. The average number of AADC mRNA copies/μm^2^ was multiplied by 10^3^ (**d**). \**P* < 0.05, and \*\*\**P* < 0.001. Data are presented as the means ±s.e.m. Details on the statistical analyses and sample sizes are provided in Supplementary Table 2.

**Extended Data Figure 5: AADC knockdown in the LHb did not change locomotion or anxiety outcomes. a–c,** OFT results: distance moved (**a**), number of centre visits (**b**), and duration spent in the centre (**c**). **d–f**, EZM results: distance moved (**d**), number of entries into open quadrants (**e**), and duration spent in open quadrants (**f**). No significant differences were detected. Data are presented as the means ±s.e.m. Details on the statistical analyses and sample sizes are provided in Supplementary Table 2.

**Extended Data Figure 6: AADC knockdown and chemogenetic stimulation of LHb D-neurons altered c-Fos expression in RMTg GABAergic neurons. a**, **b**, LHb AADC knockdown increased c-Fos expression in GAD1-positive RMTg neurons. **c**, **d**, Chemogenetic stimulation of hM3Dq-expressing LHb D-neurons decreased c-Fos expression in the GAD1-positive neurons of the RMTg. Aq, aqueduct. SERT, serotonin transporter.

**Extended Data Figure 7: Raw frequency data corresponding to Figure 5. a–d**, Firing frequencies of neurons in control mice (**a**) and AADC-knockdown mice (**b**) measured in the ON and OFF states of blue light photoactivation. Firing frequencies of neurons in control mice (**c**) and AADC-knockdown mice (**d**) measured after the application of RO5263397 (500 nM) or EPPTB (1 μM). \**P* < 0.05, \*\**P* < 0.01, and \*\*\**P* < 0.001. Data are presented as the means ±s.e.m. Details on the statistical analyses and sample sizes are provided in Supplementary Table 2.

**Extended Data Figure 8: Validation of TAAR1 mRNA expression in RMTg GABAergic neurons and LHb D-neurons. a–d**, RMTg GABAergic neurons express TAAR1 but not SERT, VGLUT2, or VGLUT3. Representative FISH images for GAD1/GAD2/TAAR1 (**a**), GAD1/GAD2/SERT (**b**), GAD1/VGLUT2/SERT (**c**), and GAD1/VGLUT3/SERT in the RMTg (**d**). **e**, Pie charts depicting the percentage of GAD1-expressing cells coexpressing other molecular markers, namely, GAD2, VGLUT2, and VGLUT3. **f**, **g**, LHb D-neurons do not express TAAR1. Representative images of FISH for TAAR1/AADC (**f**) and a pie chart depicting the percentage of TAAR1-expressing cells among AADC-positive cells (**g**). Details on the sample sizes are provided in Supplementary Table 2.

**Extended Data Figure 9: Schematic illustration of the suggested role of LHb D-neurons.**LHb D-neurons project to RMTg GABAergic neurons and are activated by acute stress. Acute stress evokes trace amine release from LHb D-neurons to RMTg GABAergic neurons, leading to neuronal inactivation via TAAR1-mediated trace aminergic signalling. Since the effect of trace amine overrides the excitatory glutamatergic transmission, LHb D-neurons act as the negative regulator of RMTg GABAergic neurons. This action ultimately activates VTA dopaminergic neurons to promote dopamine secretion into the NAc. In a depressed brain induced by chronic stress, the weakened trace aminergic signalling suppresses the activity of VTA dopaminergic neurons.

## References

1 Baik, J. H. Stress and the dopaminergic reward system. Exp Mol Med 52, 1879–1890, doi:10.1038/s12276-020-00532-4 (2020).

2 Hikosaka, O. The habenula: from stress evasion to value-based decision-making. Nat Rev Neurosci 11, 503–513, doi:10.1038/nrn2866 (2010).

3 Matsumoto, M. & Hikosaka, O. Lateral habenula as a source of negative reward signals in dopamine neurons. Nature 447, 1111–1115, doi:10.1038/nature05860 (2007).

4 Hong, S., Jhou, T. C., Smith, M., Saleem, K. S. & Hikosaka, O. Negative reward signals from the lateral habenula to dopamine neurons are mediated by rostromedial tegmental nucleus in primates. The Journal of neuroscience: the official journal of the Society for Neuroscience 31, 11457–11471, doi:10.1523/JNEUROSCI.1384-11.2011 (2011).

5 Proulx, C. D., Hikosaka, O. & Malinow, R. Reward processing by the lateral habenula in normal and depressive behaviors. Nat Neurosci 17, 1146–1152, doi:10.1038/nn.3779 (2014).

6 Shabel, S. J., Proulx, C. D., Piriz, J. & Malinow, R. Mood regulation. GABA/glutamate co-release controls habenula output and is modified by antidepressant treatment. Science 345, 1494–1498, doi:10.1126/science.1250469 (2014).

7 Li, K. et al. betaCaMKII in lateral habenula mediates core symptoms of depression. Science 341, 1016–1020, doi:10.1126/science.1240729 (2013).

8 Yang, Y., Wang, H., Hu, J. & Hu, H. Lateral habenula in the pathophysiology of depression. Curr Opin Neurobiol 48, 90–96, doi:10.1016/j.conb.2017.10.024 (2018).

9 Cui, Y. et al. Astroglial Kir4.1 in the lateral habenula drives neuronal bursts in depression. Nature 554, 323–327, doi:10.1038/nature25752 (2018).

10 Yang, Y. et al. Ketamine blocks bursting in the lateral habenula to rapidly relieve depression. Nature 554, 317–322, doi:10.1038/nature25509 (2018).

11 Holly, E. N. & Miczek, K. A. Ventral tegmental area dopamine revisited: effects of acute and repeated stress. Psychopharmacology 233, 163–186, doi:10.1007/s00213-015-4151-3 (2016).

12 Stelly, C. E. et al. Pattern of dopamine signaling during aversive events predicts active avoidance learning. Proc Natl Acad Sci U S A 116, 13641–13650, doi:10.1073/pnas.1904249116 (2019).

13 Lecca, S. et al. Aversive stimuli drive hypothalamus-to-habenula excitation to promote escape behavior. Elife 6,doi:10.7554/eLife.30697 (2017).

14 Hiraoka, K. et al. Pattern of c-Fos expression induced by tail suspension test in the mouse brain. Heliyon 3, e00316, doi:10.1016/j.heliyon.2017.e00316 (2017).

15 Jaeger, C. B. et al. Some neurons of the rat central nervous system contain aromatic-L-amino-acid decarboxylase but not monoamines. Science 219, 1233–1235, doi:10.1126/science.6131537 (1983).

16 Russo, S. J. & Nestler, E. J. The brain reward circuitry in mood disorders. Nat Rev Neurosci 14, 609–625, doi:10.1038/nrn3381 (2013).

17 Bourdy, R. & Barrot, M. A new control center for dopaminergic systems: pulling the VTA by the tail. Trends Neurosci 35, 681–690, doi:10.1016/j.tins.2012.06.007 (2012).

18 Barrot, M. et al. Braking dopamine systems: a new GABA master structure for mesolimbic and nigrostriatal functions. The Journal of neuroscience: the official journal of the Society for Neuroscience 32, 14094–14101, doi:10.1523/JNEUROSCI.3370-12.2012 (2012).

19 Yetnikoff, L., Cheng, A. Y., Lavezzi, H. N., Parsley, K. P. & Zahm, D. S. Sources of input to the rostromedial tegmental nucleus, ventral tegmental area, and lateral habenula compared: A study in rat. J Comp Neurol 523, 2426–2456, doi:10.1002/cne.23797 (2015).

20 Jaeger, C. B. et al. Aromatic L-amino acid decarboxylase in the rat brain: immunocytochemical localization in neurons of the brain stem. Neuroscience 11, 691–713, doi:10.1016/0306-4522(84)90053-8 (1984).

21 Revel, F. G. et al. TAAR1 activation modulates monoaminergic neurotransmission, preventing hyperdopaminergic and hypoglutamatergic activity. Proc Natl Acad Sci USA 108, 8485–8490, doi:10.1073/pnas.1103029108 (2011).

22 Bradaia, A. et al. The selective antagonist EPPTB reveals TAAR1-mediated regulatory mechanisms in dopaminergic neurons of the mesolimbic system. Proc Natl Acad Sci U S A 106, 20081–20086, doi:10.1073/pnas.0906522106 (2009).

23 Lindemann, L. et al. Trace amine-associated receptor 1 modulates dopaminergic activity. The Journal of pharmacology and experimental therapeutics 324, 948–956, doi:10.1124/jpet.107.132647 (2008).

24 Zhang, L., Hernandez, V. S., Vazquez-Juarez, E., Chay, F. K. & Barrio, R. A. Thirst Is Associated with Suppression of Habenula Output and Active Stress Coping: Is there a Role for a Non-canonical Vasopressin-Glutamate Pathway? Front Neural Circuits 10, 13, doi:10.3389/fncir.2016.00013 (2016).

25 Zhang, L. et al. A GABAergic cell type in the lateral habenula links hypothalamic homeostatic and midbrain motivation circuits with sex steroid signaling. Transl Psychiatry 8, 50, doi:10.1038/s41398-018-0099-5 (2018).

26 Quina, L. A., Walker, A., Morton, G., Han, V. & Turner, E. E. GAD2 Expression Defines a Class of Excitatory Lateral Habenula Neurons in Mice that Project to the Raphe and Pontine Tegmentum. eNeuro 7,doi:10.1523/ENEURO.0527-19.2020 (2020).

27 Flanigan, M. E. et al. Orexin signaling in GABAergic lateral habenula neurons modulates aggressive behavior in male mice. Nat Neurosci 23, 638–650, doi:10.1038/s41593-020-0617-7 (2020).

28 Webster, J. F. et al. Disentangling neuronal inhibition and inhibitory pathways in the lateral habenula. Scientific reports 10, 8490, doi:10.1038/s41598-020-65349-7 (2020).

29 Warden, M. R. et al. A prefrontal cortex-brainstem neuronal projection that controls response to behavioural challenge. Nature 492, 428–432, doi:10.1038/nature11617 (2012).

30 Lazaridis, I. et al. A hypothalamus-habenula circuit controls aversion. Molecular psychiatry 24, 1351–1368, doi:10.1038/s41380-019-0369-5 (2019).

31 Stamatakis, A. M. et al. Lateral Hypothalamic Area Glutamatergic Neurons and Their Projections to the Lateral Habenula Regulate Feeding and Reward. The Journal of neuroscience: the official journal of the Society for Neuroscience 36, 302–311, doi:10.1523/JNEUROSCI.1202-15.2016 (2016).

32 Ikemoto, K., Nishimura, A., Oda, T., Nagatsu, I. & Nishi, K. Number of striatal D-neurons is reduced in autopsy brains of schizophrenics. Leg Med (Tokyo) 5 Suppl 1, S221–224, doi:10.1016/s1344-6223(02)00117-7 (2003).

33 Sandler, M. et al. Deficient production of tyramine and octopamine in cases of depression. Nature 278, 357–358, doi:10.1038/278357a0 (1979).

34 Borowsky, B. et al. Trace amines: identification of a family of mammalian G protein-coupled receptors. Proc Natl Acad Sci U S A 98, 8966–8971, doi:10.1073/pnas.151105198 (2001).

35 Zucchi, R., Chiellini, G., Scanlan, T. S. & Grandy, D. K. Trace amine-associated receptors and their ligands. British journal of pharmacology 149, 967–978, doi:10.1038/sj.bjp.0706948 (2006).

36 Lindemann, L. & Hoener, M. C. A renaissance in trace amines inspired by a novel GPCR family. Trends in pharmacological sciences 26, 274–281, doi:10.1016/j.tips.2005.03.007 (2005).

37 Dodd, S. et al. Trace Amine-Associated Receptor 1 (TAAR1): A new drug target for psychiatry? Neuroscience and biobehavioral reviews 120, 537–541, doi:10.1016/j.neubiorev.2020.09.028 (2021).

38 Berry, M. D., Shitut, M. R., Almousa, A., Alcorn, J. & Tomberli, B. Membrane permeability of trace amines: evidence for a regulated, activity-dependent, nonexocytotic, synaptic release. Synapse 67, 656–667, doi:10.1002/syn.21670 (2013).

39 Lindemann, L. et al. Trace amine-associated receptors form structurally and functionally distinct subfamilies of novel G protein-coupled receptors. Genomics 85, 372–385, doi:10.1016/j.ygeno.2004.11.010 (2005).

40 Kong, Q., Zhang, H., Wang, M., Zhang, J. & Zhang, Y. The TAAR1 inhibitor EPPTB suppresses neuronal excitability and seizure activity in mice. Brain Res Bull 171, 142–149, doi:10.1016/j.brainresbull.2021.03.018 (2021).

41 Vong, L. et al. Leptin action on GABAergic neurons prevents obesity and reduces inhibitory tone to POMC neurons. Neuron 71, 142–154, doi:10.1016/j.neuron.2011.05.028 (2011).

42 Madisen, L. et al. A robust and high-throughput Cre reporting and characterization system for the whole mouse brain. Nat Neurosci 13, 133–140, doi:10.1038/nn.2467 (2010).

43 Yang, E. et al. Three-Dimensional Analysis of Mouse Habenula Subnuclei Reveals Reduced Volume and Gene Expression in the Lipopolysaccharide-mediated Depression Model. Exp Neurobiol 28, 709–719, doi:10.5607/en.2019.28.6.709 (2019).

44 Jolly, S. et al. Single-Cell Quantification of mRNA Expression in The Human Brain. Scientific reports 9, 12353, doi:10.1038/s41598-019-48787-w (2019).

45 Han, S. et al. Down-regulation of cholinergic signaling in the habenula induces anhedonia-like behavior. Scientific reports 7, 900, doi:10.1038/s41598-017-01088-6 (2017).

46 Kim, J. Y., Yang, S. H., Kwon, J., Lee, H. W. & Kim, H. Mice subjected to uncontrollable electric shocks show depression-like behaviors irrespective of their state of helplessness. Behav Brain Res 322, 138–144, doi:10.1016/j.bbr.2017.01.008 (2017).

47 Livak, K. J. & Schmittgen, T. D. Analysis of relative gene expression data using real-time quantitative PCR and the 2(-Delta Delta C(T)) Method. Methods 25, 402–408, doi:10.1006/meth.2001.1262 (2001).

48 Seo, J. S., Zhong, P., Liu, A., Yan, Z. & Greengard, P. Elevation of pll in lateral habenula mediates depression-like behavior. Molecular psychiatry 23, 1113–1119, doi:10.1038/mp.2017.96 (2018).

49 Ikemoto, K. in Trace Amines and Neurological Disorders (eds Tahira Farooqui & Akhlaq A. Farooqui) 295–307 (Academic Press, 2016).

